# Filament Formation by ChlI Challenges the Current View of Magnesium Chelatase Architecture

**DOI:** 10.64898/2026.03.06.710059

**Authors:** Natalia Łata, Łukasz Hałys, Przemysław Sendorek, Sebastian Pintscher, Paulina Indyka, Michał Rawski, Michał Gabruk

**Affiliations:** Department of Plant Physiology and Biochemistry, Faculty of Biochemistry, Biophysics and Biotechnology, Jagiellonian University, Gronostajowa 7, 30-387 Kraków, Poland; RYVU Therapeutics. Sternbacha 2, 30-394 Kraków, Poland; Doctoral School of Exact and Natural Sciences, Jagiellonian University, Łojasiewicza 11, 30-348, Kraków, Poland; Faculty of Biochemistry, Biophysics and Biotechnology, Department of Plant Biotechnology, Jagiellonian University, Gronostajowa 7, 30-387 Kraków, Poland; SOLARIS National Synchrotron Radiation Centre, Jagiellonian University, Krakow, Poland

## Abstract

Magnesium chelatase (MgCh) catalyzes the first committed step in chlorophyll biosynthesis by inserting Mg^2+^ into the tetrapyrrole ring. The enzyme comprises three core subunits: ChlI, ChlD, and ChlH, and requires the auxiliary factor GUN4 for full activity. Despite extensive investigation, the structural organization of the active holoenzyme and the mechanistic coupling between ATP hydrolysis and Mg^2+^ insertion remain poorly defined.

Here, we used cryo-electron microscopy to investigate MgCh architecture. We show that both cyanobacterial and plant ChlI homologs assemble into filamentous helical structures in the presence of Mg^2+^ and either ATP or ADP. However, only ATP-induced oligomers are susceptible to disassembly by ChlD. Structural analysis of the ATP-driven assemblies reveals compact inter-subunit packing, with ADP and Mg^2+^ coordinated at the interfacial regions. These findings suggest that ATP hydrolysis promotes subunit compaction and may facilitate partial dehydration of the Mg^2+^ hydration shell. Low-resolution reconstructions further provide a structural framework for ChlD engagement with ChlI filaments. Finally, we demonstrate that phosphatidylglycerol enhances catalytic activity, supporting a role for membrane lipids in modulating MgCh function.

**Significance statement:** Magnesium chelatase catalyzes the first committed step of chlorophyll biosynthesis, yet the structural basis of its activation and coupling to ATP hydrolysis remains unclear. Using cryo-electron microscopy, we show that the ATPase subunit ChlI forms filamentous helical oligomers in the presence of Mg^2+^ and nucleotide, a property conserved across cyanobacterial and plant homologs but not previously recognized. Only oligomers generated through ATP hydrolysis interact efficiently with the ChlD subunit, indicating that hydrolysis produces a distinct, recognition-competent conformation. Structural analysis further suggests that ATP hydrolysis promotes subunit compaction and partial dehydration of Mg^2+^. Together, these findings reveal a previously unrecognized oligomeric state of ChlI and provide a structural framework for understanding how ATP hydrolysis regulates magnesium chelatase activity.

## Introduction

Magnesium chelatase (MgCh) catalyzes the first committed step of chlorophyll biosynthesis: the insertion of Mg^2+^ into the protoporphyrin IX (PPIX) macrocycle. This entry point into the chlorophyll branch of tetrapyrrole metabolism represents a major regulatory checkpoint for chloroplast development and greening^1,2^. In contrast to ferrochelatase, which inserts Fe^2+^ into PPIX through a thermodynamically favorable reaction using a single polypeptide, Mg^2+^ chelation is energetically demanding and requires ATP hydrolysis and the coordinated activity of three subunits: ChlI, ChlD, and ChlH^3,4^. They are conserved across oxygenic photosynthetic organisms, including cyanobacteria and plants. In plant systems, however, ChlI is present in two isoforms, which have been shown to differ in their affinity towards ATP^5^. An additional regulatory element is GUN4, an accessory protein that binds porphyrins and strongly enhances MgCh activity, likely by facilitating PPIX delivery to ChlH^6^. Growing evidence indicates that GUN4, and MgCh subunits associate with plastid membranes, suggesting that membrane proximity may modulate substrate availability, protect reactive intermediates, and spatially organize porphyrin flow^7–9^.

Despite decades of research combining biochemistry, genetics, and structural studies, the overall architecture of MgCh holoenzyme and the molecular mechanism of Mg^2+^ insertion remain unresolved. Nevertheless, important mechanistic principles have been uncovered.

ChlI is an ATP-dependent oligomeric motor protein. It assembles into hexameric or semi-ring-like structures in the presence of Mg^2+^ and ATP^10,11^, while ATP hydrolysis drives conformational transitions associated with catalytic progression^3,12^. Intriguingly, multiple ATP molecules are consumed per single Mg^2+^ insertion event, indicating that ATP hydrolysis powers multiple mechanical substeps rather than one single chemical transformation^13^.

ChlD consists of an N-terminal AAA^+^-like region, a central linker, and a C-terminal integrin-like domain. Strong biochemical evidence supports the view that ChlD functions as a structural and functional bridge between ChlI and ChlH: the integrin-like C-terminal region interacts directly with ChlH^14^, while ChlD co-purifies with both ChlI and ChlH^14^. Moreover, low-resolution cryo-EM reconstructions suggest that ChlI and ChlD assemble into ring-like or pseudo-hexameric complexes whose architectures shift depending on the nucleotide state^15,16^.

ChlH is the catalytic subunit responsible for Mg^2+^ incorporation into PPIX. Its porphyrin-binding pocket has been identified through mutagenesis (Sirijovski 2008), and membrane association has been linked to substrate binding and catalytic efficiency (Yoshihara 2024). Importantly, ChlH alone is catalytically inactive, and efficient Mg^2+^ chelation strictly depends on its functional coupling to the ATP-driven ChlI–ChlD motor module^17^.

Together, these observations led to a long-standing hypothesis that MgCh forms a transient, ATP-dependent holocomplex involving multiple copies of ChlI, ChlD, and ChlH that assemble into a catalytically competent organization and dissociate after turnover^18,19^. Despite this progress, fundamental mechanistic questions remain open. The stoichiometry and lifetime of the holoenzyme complex are unknown; the temporal order of binding and release events has not been determined; the nature of ATP-coupled conformational transitions is unclear; and the mechanistic contribution of membranes has yet to be defined.

Here, by combining structural and mechanistic approaches, we examine the structural organization of MgCh. Our results challenge some assumptions underlying current models of Mg^2+^ chelation and revise the mechanistic framework through which ATP-driven motor activity is coupled to catalytic metal insertion.

## Results

After optimizing protein expression and purification, we successfully reconstituted MgCh activity in vitro and monitored the reaction by fluorescence (Fig. 1AB). Using this setup, we quantified reaction kinetics as a function of ATP or PPIX concentration under different conditions: in the presence of NADP^+^ or NADPH, and upon addition of PG, DGDG, or a thylakoid-mimicking lipid mixture (50 mol% MGDG, 35 mol% DGDG, 15 mol% PG; hereafter: OPT lipids). Control reactions lacked both lipids and dinucleotides. We found that PG consistently increased the Vmax parameter in both assay modes, whether ATP or PPIX concentration was varied (Fig. 1CD). In addition, dinucleotides produced a modest increase in Vmax, but only when ATP concentration served as the variable component (Fig. 1CD).

**Figure 1.**
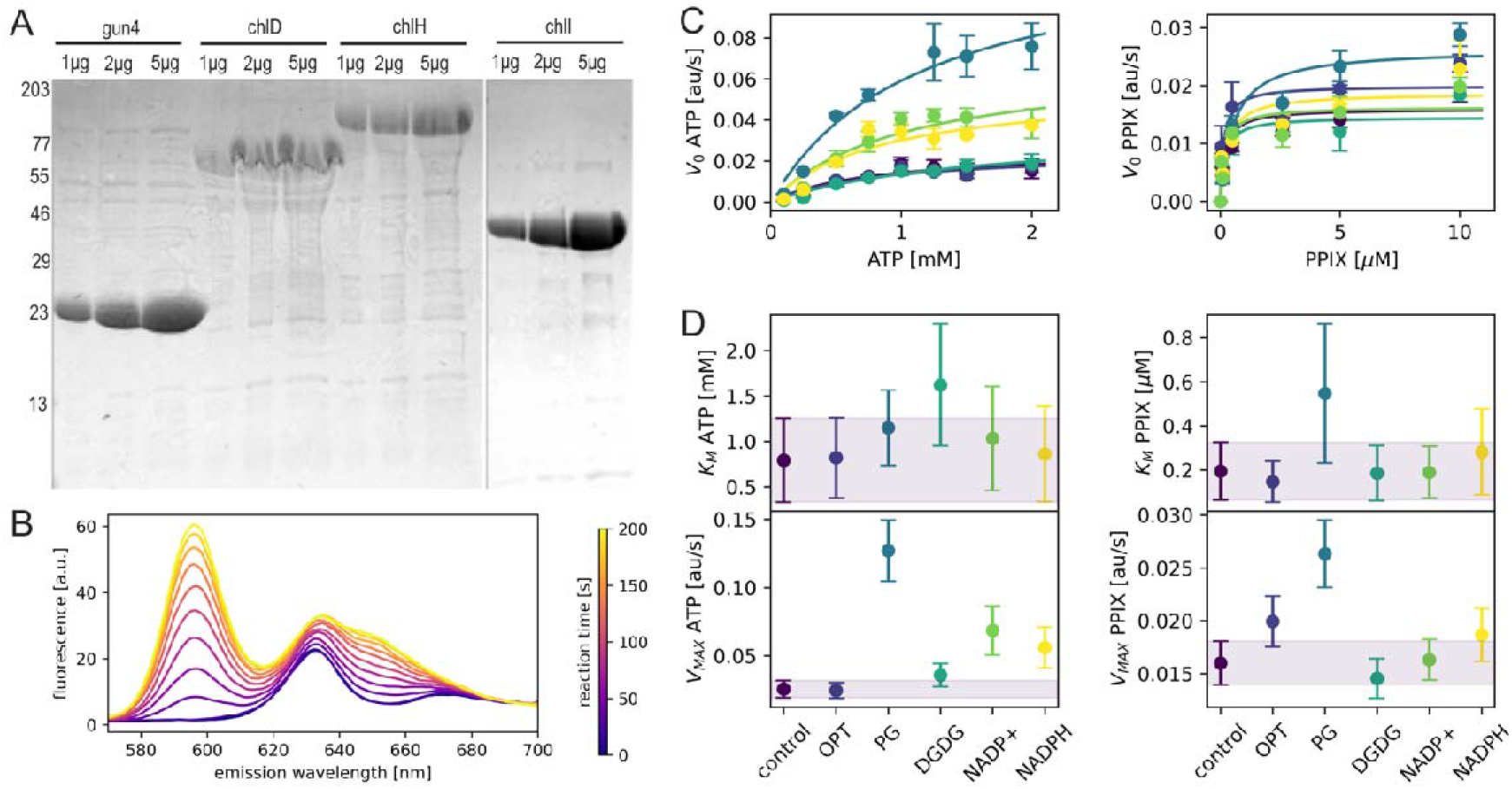
MgCh activity and its regulation. **A**. SDS–PAGE analysis of purified MgCh subunits (ChlI, ChlD, ChlH) and the auxiliary protein GUN4. **B**. Representative fluorescence spectra illustrating the progress of the MgCh reaction after mixing all subunits in the presence of Mg^2+^ and ATP. **CD**. Initial reaction velocity V_0_ (C) and apparent kinetic parameters K_M_ and V_max_ (D) of MgCh determined at varying concentrations of ATP and protoporphyrin IX (PPIX), under different conditions: with lipids PG, DGDG or a mixture resembling thylakoid membranes (OPT lipids, 50mol% MGDG, 35mol% DGDG, 15mol% PG), NADP+, NADPH, or without any additions (control conditions). The concentrations were: 0,216 μM ChlH; 0,216 μM GUN4; 1,3 μM ChlI; 1,3 μM ChD, 6 μM lipids, 200 μM dinucleotides.

Interestingly, during sample preparation, we noticed that the addition of ATP or ADP and magnesium ions induced the formation of a white turbidity in the ChlI solution within a few minutes (Fig. 2A). Electron microscopy revealed that this material consisted of filamentous assemblies (Fig 2B). To investigate the oligomerization kinetics, we took advantage of the turbidity formation and monitored scattering. We found that ADP promoted the oligomerization at much lower concentrations comparing to ATP, however, only the ATP-induced oligomers were successfully disassembled by the addition of ChlD (Fig. 2CD).

**Figure 2.**
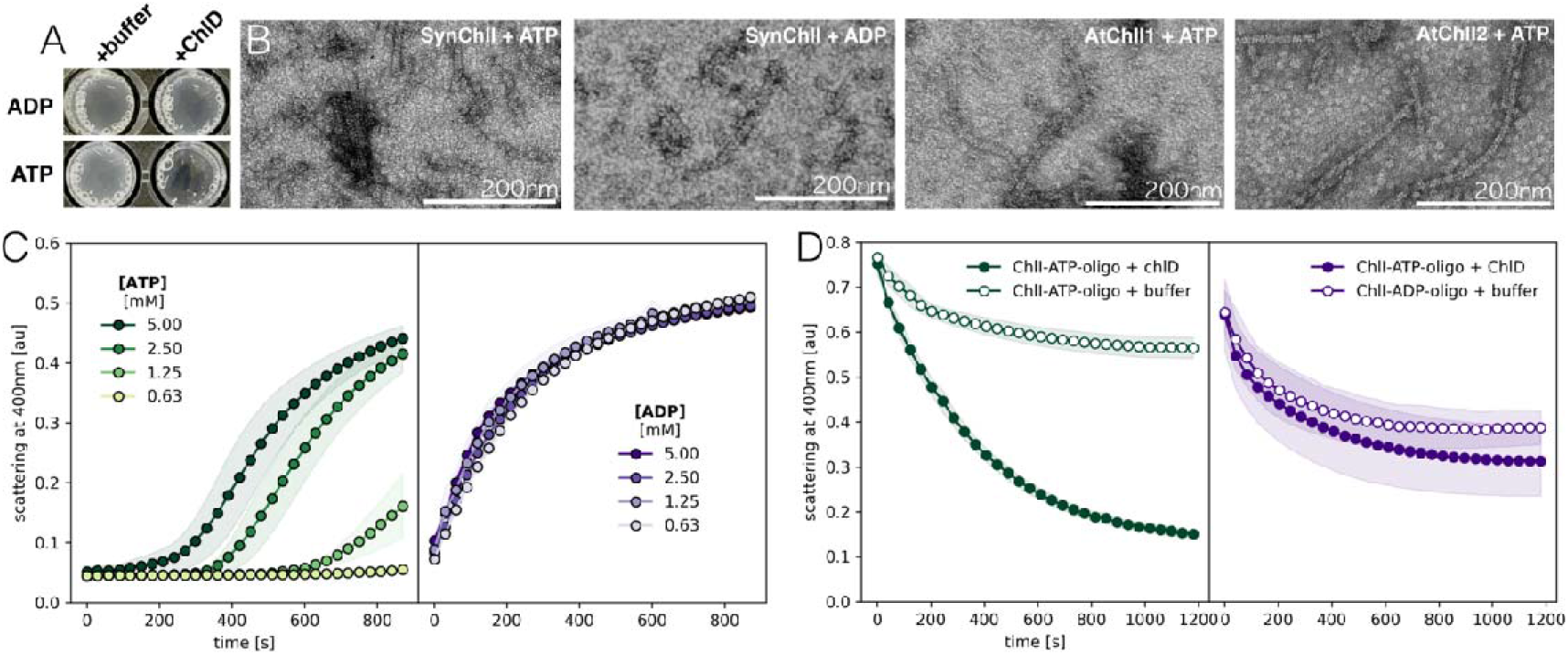
ChlI form filamentous oligomers in both Synechocystis and Arabidopsis. Representative negative-stain electron microscopy micrographs showing individual filaments and filament aggregates.

To determine whether this behavior is conserved across ChlI proteins, we examined the two *A. thaliana* isoforms, ChlI1 and ChlI2 (Fig 2B). Both formed similar filaments, indicating that filamentous self-assembly is an intrinsic property of the I subunit of MgCh. This finding prompted us to investigate the structural organization of the filaments by cryo-EM.

During the screening of the cryo-grids, we noticed that the filaments of chlI from Synechocystis produced with Mg2+ and ATP, were too clumped and were too dense. Therefore we used slowly hydrolyzable ATP analog, tetralithium salt of adenosine 5′-O-(3-thiotriphosphate), hereafter aATP, that’s still initiated filament formation but at smaller scale.

We collected 11863 movies, 40 frames each, for ChlI:aATP:Mg^2+^ sample that allowed us to reconstruct the map at 2.94 Å resolution, which revealed a helical nature of the assembly (Fig. 3ABC). Based on the map, we built a model comprising seven subunits and thanks to the stabilization of the subunits between the turns of the helix, we were able to model the full-length amino acids chain, including the two flexible loops (92-101, 125-136), that were unstructured in previous models (Fig. 3D).

**Figure 3.**
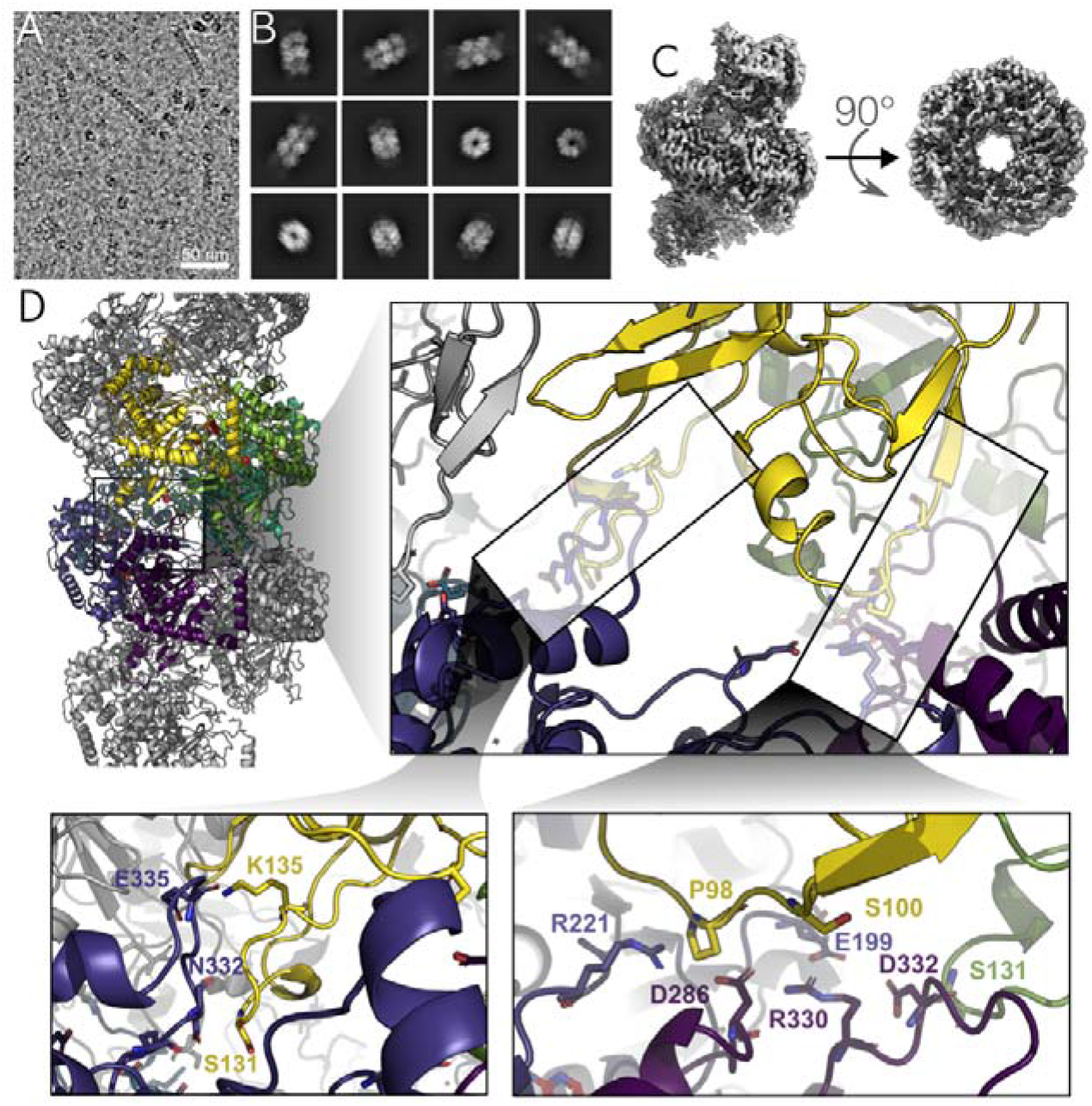
Structure of the ChlI oligomer. **A**. Cryo-EM micrograph showing a visible ChlI filament. **B**. Representative 2D class averages. **C**. Cryo-EM 3D reconstruction of ChlI oligomer. **D**. Model of ChlI oligomer with close-up views of inter-oligomer interactions between subunits.

Interestingly, our model revealed tight interactions between adjacent subunits at the nucleotide-binding interfaces, where adenosine nucleotides and Mg^2+^ ions are coordinated. Despite the use of slowly hydrolysable ATP analog, all of the molecules we identified were ADP (Fig. 4AB), what indicates that the nucleotides underwent hydrolysis triggered by the ChlI. The model reviled that multiple residues extend toward the magnesium ion, forming a dense interaction network (Fig. 4CD). Comparison with previously reported structures showed that the distance between neighboring subunits, measured as the Cα–Cα distance between residues D153 and E206, was smallest in our assembly (Fig. 4F). In contrast, other structures displayed larger and more heterogeneous spacing (Fig. 4EF). The largest separation was observed in the nucleotide-free crystal structure (PDB: 6L8D). Smaller but still variable distances characterized the hexameric assemblies (PDB: 8OSF and 8OSH), which contain five or four bound ATP molecules, respectively. A pentameric assembly displayed even shorter average distances, although its low resolution precluded determination of the ATP/ADP ratio. Among all available ChlI oligomer structures, the model presented here exhibits the shortest and least dispersed inter-subunit distances, with ATP hydrolyzed in each binding site (Fig. 4ED). These observations lead us to speculate that ATP hydrolysis may promote conformational changes that result in subunit compaction, potentially by facilitating dehydration of the magnesium ion hydration shell. However, because this putative role of ATP, together with the previously unreported oligomeric state of ChlI presented in this work, is difficult to reconcile with current mechanistic models of MgCh, we prepared three additional cryo-EM sample, containing ChlI and ChlD in the presence of ATP and Mg^2+^. Although the resulting dataset (10386 movies) did not yield high-resolution reconstructions due to severe preferred-orientation effects, the 2D class averages and low-resolution maps revealed several unexpected features (Fig. S1).

**Figure 4.**
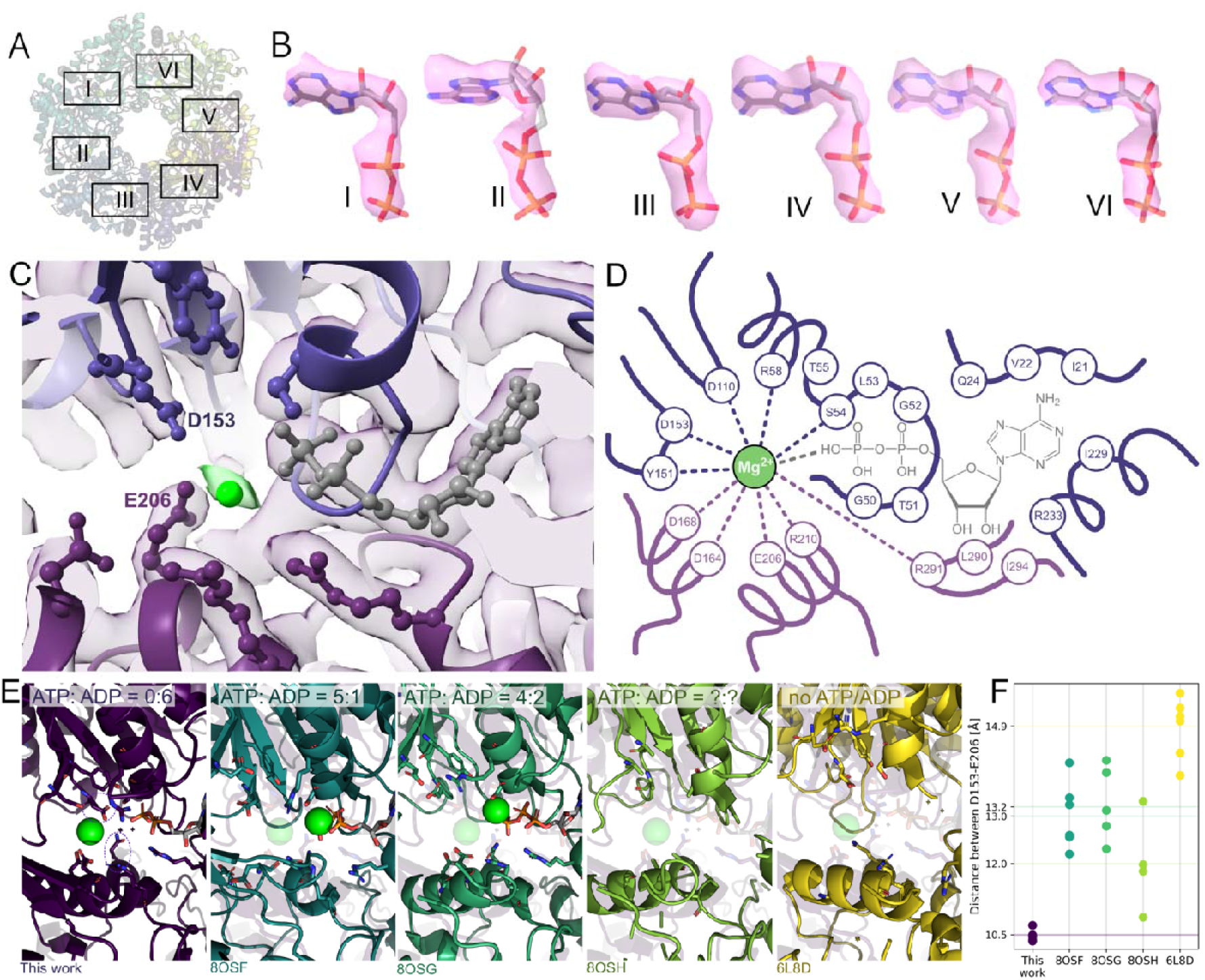
Nucleotide-binding site in ChlI. **A**. Top-down view of the ChlI oligomer model with six oligomerization interfaces indicated. **D**. Densities corresponding to nucleotides located at the oligomerization interfaces highlighted in panel A. **C**. Close-up view of the nucleotide-binding site with map density visible. **D**. Schematic representation of residues involved in nucleotide binding at the oligomerization interface between two ChlI subunits. **E**. Comparison of the nucleotide-binding site observed in the structure determined in this study with previously reported models. The semi-transparent structure in the background represents the model determined in this study. All models are aligned to the subunit shown in the lower part of the images. Residues D153 and E206 are indicated by dotted circles. **F**. Inter-subunit distances within the oligomer, calculated as distances between the Cα atoms of residues D153 and E206 in adjacent subunits.

In the ChlI-ChlD sample, oligomeric ChlI filaments were notably absent. Instead, we observed ChlD monomers and semiring-like oligomers resembling ChlI pentamers. Closer inspection revealed additional density at the open ends of these semirings, consistent in size and shape with the N-terminal domain of ChlD. This confirms that ChlD can bind to and remodel ChlI oligomers (Fig. S1).

## Discussion

### Regulation of MgCh activity

In our study, we observed a positive effect of PG on Vmax when either ATP or PPIX was titrated. The fact that PG increases Vmax under both conditions suggests that anionic lipids enhance a step common to ATP- and PPIX-dependent phases of the reaction. Mechanistically, this may reflect enhanced PPIX delivery by GUN4 or more efficient communication between ChlD and ChlH during catalytic turnover.

These interpretations align with earlier findings that both ChlD and ChlH can associate with membrane lipids in both plants and cyanobacteria^14,20,21^, with ChlH showing membrane binding in a Mg^2+^-dependent manner^22^. Moreover, the physiological relevance of PG for MgCh has been indicated previously by the phenotype of etiolated seedlings partially deficient in PG biosynthesis^23^. Together, these observations highlight a potentially important role for PG in modulating MgCh activity in vivo. However, a definitive mechanistic understanding will require structural information on ChlH embedded within or interacting with a lipid membrane.

The stimulatory effect of NADP(H) on Vmax in ATP-dependent assays is unexpected. Earlier studies showed that ChlI ATPase activity can be enhanced indirectly through thioredoxin^24^, which relies on NADPH, but no direct influence of NADP(H) on MgCh has been reported. The underlying mechanism thus remains unclear and requires further investigation.

### ChlI oligomerization

Our data demonstrate that ChlI forms filamentous oligomers in the presence of Mg^2+^ and either ADP or ATP. That property of I subunit seems to be conserved across both *Arabidopsis* isoforms and the *Synechocystis* homolog, but surprisingly this behavior has not been reported previously. Crucially, our filament reconstruction resolves the full-length ChlI polypeptide, including loops that are unmodelled in previous structures^10,11^ and which form contacts across successive helical turns. These features strongly support the interpretation that the observed assembly reflects a physiologically relevant oligomeric state rather than an artefact.

Although ATP- and ADP-induced assemblies converge to an ADP–Mg^2+^-bound state, our data suggest that the oligomers generated via ATP hydrolysis represent a structurally distinct population. Notably, only assemblies formed in the presence of ATP display efficient interaction with ChlD, indicating that hydrolysis is coupled to the formation of a recognition-competent conformation. We speculate that ATP hydrolysis drives a conformational rearrangement, potentially involving alterations in helical pitch and/or surface residue exposure, selectively recognized by the D subunit. Thus, ATP hydrolysis may imprint structural information that is not evident from nucleotide occupancy alone.

Consistent with this interpretation, our analysis shows that ChlI filaments contain exclusively hydrolyzed nucleotide, despite earlier studies predominantly reporting ATP-bound^10,11^. Based on the structural compaction observed within the ChlI filament, we hypothesize that ATP hydrolysis contributes to both subunit tightening and partial dehydration of Mg^2+^. These observations align with the nucleotide-free crystal structure, where ChlI adopts a planar, more relaxed ring conformation^10^.

## Role of ChlD

Although our semi-ring reconstruction is of modest resolution, the ChlI architecture is unambiguously identifiable. Additional densities at the open ends of the assembly can accommodate the N-terminal domain of ChlD. Given that this region belongs to the AAA^+^ ATPase family, it is plausible that ChlD binds filament termini. This interpretation is supported by atomic force microscopy experiments, which showed that both the N-terminal segment and central polyproline region of ChlD are required for ChlI–ChlD interactions^12^.

We demonstrate that ChlD actively disassembles ChlI filaments. However, due to the limited resolution of the map, the stoichiometry of the ChlI–ChlD complex remains uncertain and could correspond to either a 2:2 or a 3:1 ratio. Previously proposed models suggested either 6:6^19,27,28^ or 2:1^25^ ratio but these values were proposed based on biochemical assays rather than structural information. Regardless of the stoichiometry, the mechanistic link between conformational rearrangements in ChlI and Mg^2+^ transfer to ChlH via ChlD remains elusive. In particular, how ChlD orchestrates Mg^2+^ loading and delivery, and how ATP-driven conformational changes in ChlI are propagated to ChlH, are still unresolved. Resolving these questions will require further high-resolution structural investigation.

## Materials and methods

### Gene cloning

Genes encoding the MgCh subunits were obtained as follows: *chlI* and *gun4* were amplified from genomic DNA of *Synechocystis sp*. PCC 6803, whereas *chlD* and *chlH* were commercially synthesized as codon-optimized sequences for heterologous expression in *E. coli*.

Amplification of the desired fragments was performed by PCR using specific primers (Table S1). The PCR reaction mixture contained: Q5 Reaction Buffer (New England Biolabs, USA), 2 μM of each primer, 0.2 mM dNTP mix, 1 ng of template DNA, and 1 U Q5 High-Fidelity DNA Polymerase (New England Biolabs, USA). Reactions were performed under conditions suggested by manufacturer of the polymerase with annealing temperature set to 68°C.

PCR products were separated on 1% agarose gels in TAE buffer. The separated products were cut out of the gel and purified with the GeneJET Gel Extraction Kit (Thermo Fisher Scientific, USA), following the manufacturer’s protocol.

Amplified fragments were ligated into linearized pET15b plasmid (amplified by PCR) using Gibson assembly^29^ with NEBuilder HiFi DNA Assembly Master Mix (New England Biolabs, USA). The molar ratio of insert to plasmid was set to 2:1. The transformant colonies were verified with colony PCR and then with sequencing.

### Protein expression and purification

Expression of MgCh subunits was carried out in *E. coli* BL21(DE3). Overnight precultures (25 mL) were used to inoculate 1 L of fresh LB medium, followed by incubation at 37 °C until OD600 reached 0.5–0.8. Protein expression was induced with 0.4 mM IPTG. Cultures were grown overnight (12 h) at 37 °C with shaking (180 rpm). Cells were harvested by centrifugation (12,000 × g, 4 °C, 4 min) and stored at −20 °C until further purification.

Cell pellets were resuspended in phosphate buffer (300 mM Na2HPO4, 50 mM NaCl, 7 mM 2-mercaptoethanol, pH7.1) and disrupted by sonication on ice-water bath using a Sonics Vibracell VC505 sonicator (6 s pulse on, 14 s off, 40% amplitude, total sonication time 9 min). The lysate was clarified by centrifugation (18,000 × g, 45 min, 4 °C).

The supernatant was incubated with 5 mL of TALON Metal Affinity Resin (Takata Bio Inc., Japan) for 2 h at 4 °C with gentle mixing. After two washes with the phosphate buffer, bound proteins were eluted in fractions by incubation with 4 mL elution buffer (the phosphate buffer enriched with 200 mM imidazole, pH7.1) for 15–5 min at 4 °C, followed by centrifugation (1,100 × g, 35 s) in AmiconPro columns (Merck KGaA, Germany).

Protein concentration was initially determined spectrophotometrically using a NanoDrop Lite instrument (Thermo Fisher Scientific, USA). Fractions containing >0.5 mg/mL protein were pooled, supplemented with glycerol to a final concentration of 25% (w/v), and quantified by the Bradford assay according to the manufacturer’s instructions (Sigma-Aldrich, USA) [36].

### Fluorescence spectroscopy

Prior to activity assays, storage buffer of purified MgCh subunits was exchanged for reaction buffer using PD midiTrap G-25 desalting columns (GE Healthcare, UK) according to the manufacturer’s instructions. Reaction buffer contained: 50 mM Tricine-NaOH (pH 8.0), 10 mM NaCl, 7 mM 2-merkaptoetanol, 15 mM MgCl_2_ and 21 mM glycerol.

Protein mixtures were prepared as follows: (i) ChlH + GUN4 + PPIX, and (ii) ChlI + ChlD + ATP (Mg2+ ions were in the buffer). Mixtures were preincubated for 15 min at 30 °C. The reaction was initiated by combining the two mixtures. Final concentrations were: 360 nM ChlH, 360 nM GUN4, 260 nM ChlI, 30 nM ChlD, 360 nM PPIX (Sigma-Aldrich, USA), and 1 mM ATP (Sigma-Aldrich, USA) [11].

Fluorescence spectra were recorded on a PerkinElmer LS 55 spectrofluorometer (PerkinElmer, USA) under the following conditions: excitation at 420 nm; emission range 500–700 nm; scan speed 500 nm/min; slit width 10 nm. In addition, fluorescence intensity at 595 nm was monitored over 500 s.

### Kinetic measurements

For kinetic measurements, proteins were buffer-exchanged as described above. Two mixtures were prepared: (i) ChlH + GUN4 + PPIX and (ii) ChlI + ChlD + ATP, which were incubated for 15 min at 30 °C, and then combined immediately before measurement. Final concentrations were: 0.216 μM ChlH, 0.216 μM GUN4, 1.3 μM ChlI, 1.3 μM ChlD, ATP (2 mM or 0.1–8.5 mM), and PPIX (2.6 μM or 0.01–10 μM).

Reactions were performed at 30 °C in the absence or presence of 6 μM lipids (PG, DGDG, or OPT lipids – a mixture of MGDG/DGDG/PG, 50/35/15 mol%, respectively), or in the presence of 200 μM nucleotides (NADPH or NADP+). The increase in product formation was monitored as fluorescence emission at 596 nm over 5 min using a FlexStation 3 Multi-Mode Microplate Reader (Molecular Devices UK Limited, UK), with acquisition every 12–24 s.

Fluorescence intensity traces obtained at different substrate concentrations were plotted against time (Figures 24–29). Initial reaction rates (V0) were determined from the slopes of tangents fitted to the first 126 s of the reaction. V0 values were plotted against substrate concentration (S), and the data were fitted to the Michaelis–Menten equation:

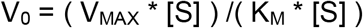

Kinetic parameters (Michaelis constant KM, and maximum velocity V_max_) and their uncertainties were derived from the fits.

### Preparation of cryo-EM sample

For the cryoEM analysis, proteins were buffer-exchanged as described above and incubated for 15 min at 30 °C. Four different mixtures were prepared: 1) ChlI with 15mM MgCl_2_ and 1mM tetralithium salt of adenosine 5′-O-(3-thiotriphosphate) (total protein concentration: 1mg/ml); 2) ChlI and ChlD with 15 mM MgCl_2_ and 2.5 mM ATP (molar ratio 1:1, total protein concentration: 1mg/ml); 3.5 µL of the mixture were applied to Quantifoil R2/1, Cu 200-mesh grids. Grids were glow-discharged immediately before use (60 s, 8 mA) and vitrified on a Vitrobot Mark IV (Thermo Fisher Scientific) under the following conditions: 100% humidity, 277 K, blot time 3 s, wait time 0 s, blot force 3, drain time 0 s, one blotting cycle. After plunge-freezing in liquid ethane, grids were stored in liquid nitrogen until use.

### cryoEM data collection, image analysis and 3D reconstructions

Twp datasets were collected at the National Cryo-EM Centre SOLARIS in Kraków, Poland using Titan Krios G3i microscope (Thermo Fisher Scientific) at an accelerating voltage of 300□kV, a magnification of ×105,000 and a pixel size of 0.86□Å using EPU 2.10.0.1941REL software. A K3 direct electron detector was used for data collection in a BioQuantum Imaging Filter (Gatan) set-up with a 20□eV slit width. The detector was operated in counting mode. The defocus range applied was −2.1□µm to −0.9□µm with 0.3□µm steps. The dataset consisted of: 1) 11863 movies with 41.38 e/A2 dose (chlI), and 2) 10386 movies with 41.38 e/A2 dose (ChlI + ChlD).

Cryo-EM movies were imported into CryoSPARC v4.7^30^ and processed using patch-based motion correction, followed by patch-based CTF estimation. Motion-corrected micrographs were calibrated with a physical pixel size (0.84 Å/pixel).

Particle picking was performed using the Topaz^31^ neural network picker based on a ResNet architecture. Initial particle picking and all early processing steps were carried out on downsampled micrographs (binning factor: 4).

Particles were extracted from micrographs using a box size (box size: 400 px) and subjected to multiple rounds of 2D classification to remove false positives and low-quality particles. Selected 2D class averages were used for ab initio 3D reconstruction, followed by heterogeneous refinement.

The output of heterogeneous refinement was used to iteratively retrain the Topaz picker. The retrained model was applied to the full dataset, picked particles were extracted and subjected to additional rounds of 2D classification, ab initio reconstruction, heterogeneous refinement, and selected volumes were further refined using non-uniform refinement, all performed on downsampled particles. Then, particles assigned to the highest-quality class were re-extracted from the original, unbinned micrographs (binning factor: 1) and subjected to non-uniform refinement to generate the final volume.

Atomic model of CHlI was built based on AlphaFold^32^ model using ChimeraX^33^ and Coot^34^, refined using PHENIX^35^ and validated with MolProbity^36^.

## Acknowledgments

This work was supported by SONATA project (2019/35/D/NZ1/00295) granted by National Science Centre (NCN) to MG. The cryo-EM study was supported under the Polish Ministry of Education and Science project: ‘Support for research and development with the use of research infrastructure of the National Synchrotron Radiation Centre SOLARIS’ under contract number 1/SOL/2021/2. Data collection was performed under proposal nr. 233013. We also acknowledge the Polish high-performance computing infrastructure PLGrid (HPC Center: ACK Cyfronet AGH) for providing computer facilities and support within computational grant numbers: PLG/2024/017733, PLG/2025/018860, and PLG/2026/019118.

## Author contributions

The study was designed and directed by M.G. Experiments were conducted by N.Ł., Ł.H. and P.S. Grids freezing and initial grid screening was performed by S.P. and M.G. Data collection was done by M.R. and P.I. Data processing was performed by M.G. and S.P. The paper was written by M.G. with all authors discussing the results and refining and approving the final version.

## Data availability

The cryo-EM densities have been deposited in the Electron Microscopy Data Bank (EMDB) and the protein model have been deposited in the Protein Data Bank (PDB) with the following accession codes: ChlI oligomer: 29BZ / EMD-57069; ChlI:ChlD putative complex: EMD-57070, putative ChlD: EMD-57071.

