## Supplementary material for "Filament Formation by ChlI Challenges the Current View of Magnesium Chelatase Architecture": S1

| 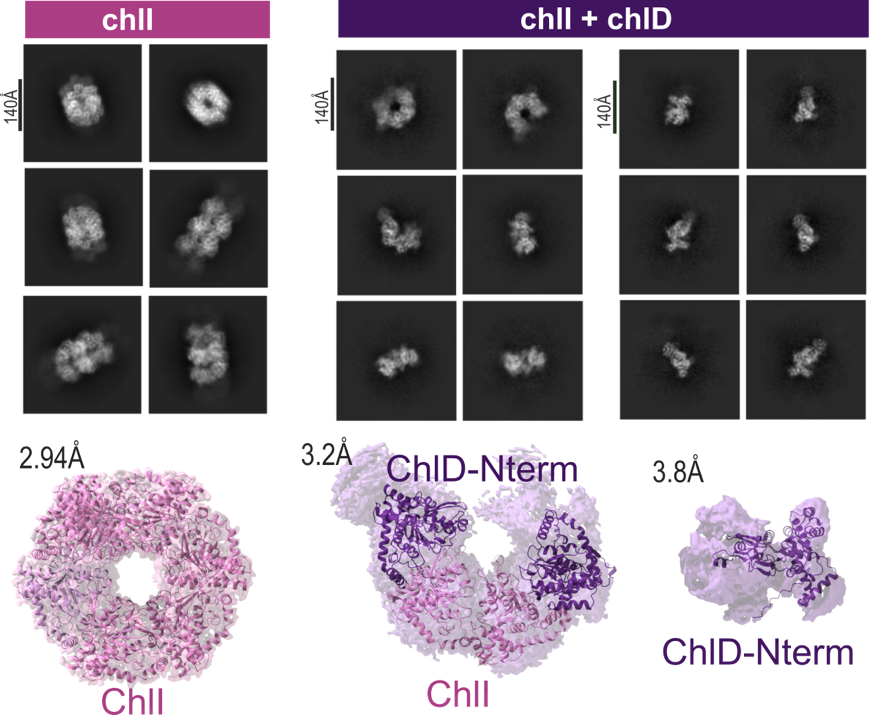 | **Figure S1. Structure of CHlI depends on the presence of ChlD.** Representative 2D class averages obtained from different mixtures and the corresponding 3D volumes. AlphaFold models of individual subunits are fitted into the volumes for size and shape comparison. |
| --- | --- |

**Table S1. Sequence of primers used for gene cloning.**

| PCR product | Sequence of primers |
| --- | --- |
| *GUN4* | F: CCGCGCGGCAGCCATATGTCTGATAATTTGACCGAACCGAAC  R: TGTTAGCAGCCGGATTACCAACCGTATTGGGAC |
| *ChlI* | F: CCGCGCGGCAGCCATATGACTGCCACCCTTGC  R: TGTTAGCAGCCGGATTAAGCTTCATCGACAACGC |
| *ChlD* | F: CCGCGCGGCAGCCATATGACAACCTTGACCCCGTTTATC  R: TTGTTAGCAGCCGGATTACTGCATATCTGCAATCGCCTG |
| *ChlH* | F: CCGCGCGGCAGCCATATGTTTACGAATGTCAAGTCGACCATTC  R: TTGTTAGCAGCCGGATTACTCTACACCCTCGATGCGATC |
| *pET15b* | F: TCCGGCTGCTAACAAAGCCCGA  R: ATGGCTGCCGCGCGGCAC |

| **Table S2. ChlI oligomer cryo-EM data collection and refinement statistics.** | |
| --- | --- |
| Data collection |  |
| Magnification | x105,000 |
| Voltage (kV) | 300 |
| Electron Microscope | FEI Titan Krios G3i |
| Defocus (um) | -0.9 to -2.7 |
| Pixel size (Å) | 0.86 |
| Total dose (e^-^/ Å^2^) | 41.38 |
| Number of frames | 40 |
| Number of micrographs | 11 863 |
| Refinement |  |
| Number of total particles | 226,517 |
| GS-FSC Resolution (0.143, Å)^a^ | 2.94 |
| Model composition |  |
| Chains | 7 |
| Protein residues | 2387 |
| r.m.s.d. |  |
| Bond lengths (Å) | 0.004 (0) |
| Bond angles (°) | 0.895 (13) |
| Validation |  |
| MolProbity score | 1.78 |
| Clash score | 8.76 |
| Ramachandran plot |  |
| Favored (%) | 96.17 |
| Allowed (%) | 3.71 |
| Disallowed (%) | 0.13 |
| CC Mask | 0.85 |
| ^a^Gold-Standard Fourier-Shell Correlation |  |
